# Influence of Diabetes Mellitus Type 2 on the angiogenic potential of Bone Marrow derived Mesenchymal Stem Cells

**DOI:** 10.1101/2023.08.29.555445

**Authors:** Sofia Adwan, Hana Hammad, Mamoun Ahram, Abdalla Awidi

## Abstract

Mesenchymal stem cells (MSCs) are used as novel therapeutic tools in cell-based therapies including diabetic complications. However, diabetes may negatively affect the therapeutic potential of autologous MSCs. In order to enhance the utilization of diabetic MSCs, better characterization of their angiogenic capacity should be performed. The aim of this study was to investigate the effect of diabetes mellitus (DM) on the angiogenic potential of bone marrow derived MSCs. MSCs were isolated from DM subjects and were compared with MSCs isolated from non-DM subjects. The angiogenic potential of MSCs was assessed in which there was insignificant difference in the proliferation and expression of angiogenic factors between the two groups. Moreover, no statistically significant change was found in the viability and angiogenic activities of endothelial cells isolated from both groups. Results indicate that the hyperglycemic milieu had no significant impact on the angiogenic-related functional properties of MSCs and they are able to survive the harsh diabetic conditions.

## Introduction

Mesenchymal stem cells (MSCs) are multipotent cells that are mounting as an ideal candidate cell source in regenerative medicine due to their unique properties such as their capacity to self-renew, easy accessibility, immunomodulatory effect, and lack of eliciting immune response [1–3]. They can be isolated from various sources where the bone marrow (BM) is the most common source of MSCs [4]. Despite the broad therapeutic application of MSCs, they form a rare population comprising of only 0.001-0.01% of the total bone marrow mononuclear cells (MNCs)[5]. Therefore, their use in clinical application requires extensive *in vitro* expansion prior to their use in therapeutic settings. Nonetheless, the expansion potential is limited since the length of expansion and quality of cells are strongly affected by patient’s health status, genetic makeup and age. This could have important implications for stem cell therapies as differences in patient’s metabolic status may have an impact on the therapeutic function of MSCs [6]. It has been described that MSCs isolated from patients with type II diabetes mellitus are characterized by increased apoptosis, autophagy, accumulation of reactive oxygen species and mitochondria degradation[7].

Stemming from the presented data, therapeutic application of MSC-based angiogenesis isolated from these patients may be limited due to their dysfunctionality. Diabetes Mellitus (DM), which is recognized as one constituent of metabolic syndrome disorders, is the most common cause of hyperglycemia [8]. Persistent exposure of the endothelium to high levels of glucose causes malfunction of the vasculature, complicating the pathogenesis of diabetes [9]. Furthermore, hyperglycemia has been demonstrated to change the hematopoietic niche of the bone marrow and impair its hematopoietic function [10]. In addition, hyper-glucose has been shown to induce premature senescence and genomic and telomeric alterations as well as alterations in chemokine expression in MSCs [11]. Thus, one of the limitations in the use of autologous MSCs as therapeutic agents in diabetic patients is the impaired paracrine angiogenic potential in addition to low transdifferentiation into endothelial cells (ECs) [12]. The principal objective of this study was to compare non-DM MSCs and DM-MSCs with respect to their growth, surface markers, differentiation capacity and their ability in inducing angiogenesis in an *in vitro* endothelial cell model.

## Materials and Methods

Ethical approval was obtained from the IRB committee of the cell therapy center and written informed consent in accordance with Helsinki declaration was obtained from all donors prior to any sample collection. Six bone marrow samples were collected from six patients, these 6 samples were divided equally into two subgroups: non-diabetic (non-DM MSCs) and diabetic (DM-MSCs). Non-DM donors had no congenital or acquired pathologies and were negative for any infectious or systemic disease. Diabetic patients had type 2 diabetes mellitus but were also negative for any infectious disease.

### Isolation of MSCs from bone marrow aspirates

Human MSCs were isolated from BM aspirates based on their plastic adherence properties and were characterized according to the International Society of Cellular Therapy (ISCT) criteria for defining MSCs [19]. Mononuclear cells (MNCs) were separated by density gradient method and were seeded at a density of 0.16-0.18x 10^6^ cells/cm^2^ in a complete α-MEM culture medium (Gibco, Thermo Fisher, USA). Cells were allowed to attach for 24 hours then the medium and nonadherent cells were removed to be replaced with fresh complete MSC medium. On daily basis, the cells were observed, and the culture medium was replaced every 3 days until individual colonies reached 70-80% confluence and became ready for subculturing.

### Surface marker characterization

Characterization of MSC surface markers was performed according to ISCT guidelines. Surface markers of MSCs were tested using BD Stemflow^TM^ hMSC Analysis kit (BD, USA).

### Growth kinetic assessment

Growth kinetics were assessed in order to examine population doubling time (PDT) and to predict the expansion yield. This was achieved through long-term culture expansion of control and diabetic bone marrow MSCs. In this assay, the cells were seeded at an initial density of 4000 cells/cm^2^ and then split after they reach 70-80% confluence, counted and reseeded to the initial seeding density of 4000 cells/cm^2^.

### Preparation of cell conditioned medium (CCM)

In order to investigate the therapeutic effect of MSCs on the angiogenesis of HUVECs, CCM was collected from the cultured bone marrow MSCs to assess the difference between the angiogenic potential of the diabetic and nondiabetic MSCs. Cells were plated at a seeding density of 4000 cells/cm^2^. When these cells reached 80-90% confluence, the spent medium was discarded and flasks were washed three times with PBS, serum free medium (SFM) was added to the cells and then incubated. After 48 hours, the CCM was collected, centrifuged at 1000 xg for 10 minutes to remove any cell debris, filtered through 0.22 µm syringe filters, aliquoted and finally stored at − 80°C until use. The CCM used was diluted with M199 to a total concentration of 1:1. For the measurement of growth factors, CCM was concentrated using 3-KD cut off filtration tubes (Millipore, USA) and stored at −80^0^C to be analyzed later [20].

### Multilineage Differentiation

Passage 4 bone marrow MSCs (non-DM and DM) were used to assess their multilineage potential toward osteogenic and adipogenic lineages. The ability of MSCs to differentiate toward osteogenic and adipogenic lineage was assessed by StemPro® Ostogenesis Differentiation and StemPro® Adipogenesis Differentiation Kits (Gibco, USA), respectively.

### Isolation of HUVECs

After the informed consent had been acquired from mother donors. Umbilical cords (UC) were obtained from normal, full-term births, after either Cesarean or normal vaginal delivery. The UC was collected and HUVECs were isolated by enzymatic digestion method described by Crampton et al., 2007 [21]. And then cells were seeded in 25-cm^2^ tissue culture flask pre-coated with 0.1% Gelatin (Sigma-Aldrich, USA). When cultures reached 70-80% confluence, reseeding was performed with a subcultivation ratio of 1:2 to 1:3. Complete M199 culture medium consisting of M199 medium (Gibco, Thermo Fisher, USA) supplemented with 10% Fetal Bovine Serum (FBS), 1% (w/v) P/S, 2 mM L-glutamine, 0.1 mg/ml Heparin (Sigma-Aldrich, USA) and 0.03 mg/ml Endothelial cell growth supplement (ECGS) from bovine neural tissue (Sigma-Aldrich, USA) was used. Cells were incubated and medium renewal was performed 2-3 times per week. At passage 2-6, cells were used for subsequent experimental work including characterization of HUVECs, migration, wound healing and tube formation assays.

### Characterization of HUVECs

The HUVECs were further characterized by flow cytometry to assess their purity using the following antibodies: CD31.PE, CD144.PerCP-Cy5.5, CD45.BB515 and CD34.BV421.

### Angiogenic growth factor analysis

The Human angiogenesis antibody array kit (R&D systems, USA) was used to detect the relative levels of 55 angiogenesis-related proteins in a single sample. The protocol was carried out as per manufacturer instructions. Corresponding signals on different arrays were compared to determine the *relative* change in angiogenesis-related proteins between different samples.

### Cell proliferation assay

An (MTT) assay was used to assess cell viability of HUVECs treated with non-DM and DM-MSC CCM along with complete M199 and SFM as controls. CellTiter Non-Radioactive Cell Proliferation MTT Assay Kit (Promega; USA) was used according to the manufacturer instructions. Cell viability was tested by MTT assay for 120 hours. HUVECs were plated overnight (day 0) and were subsequently treated with non-DM, DM-MSC CCM, complete M199 and SFM.

### Cell migration assay

The chemotactic motility of HUVECs was determined using modified Boyden chambers as described by Hwang *et al.,* (2012) [22] with some modifications. Polycarbonate membranes with 8 μm pores (Corning, USA) were used in a 24-well format. A total of 5×10^4^ HUVECs in 100 µl cell suspension, serum-starved overnight were added to the upper compartment. The lower compartment was filled with 750 µl of different media including non-DM and DM-MSC CCM, SFM (negative control) and standard cell culture medium (complete M199 as positive control). HUVECs were allowed to migrate for 24 hours, after which chambers were removed washed with PBS. Non-migrated cells were removed with a cotton swab, and then were fixed for 2 minutes in 4 % formaldehyde in PBS. Chambers were then washed twice with PBS. Membranes were stained with ProLong® Diamond antifade mountant (Invitrogen ThermoFisher, USA).

### Wound healing assay

The wound healing method was used to assess cell migration ability. It was examined as reported previously with some modification [23]. In brief, 7 x 10^5^ HUVECs were seeded onto 6-well plates. Cells were starved for at least 4 hours, and then a wound was created as a line across the well using a 200-μL pipette tip. After wounding, cell debris were removed by washing the cells with warm serum free medium. Cells that had migrated into the wound area or with extended protrusion from the border of the wound were photographed and quantified after 12 hours using an inverted microscope. The cell-free area at 12h was determined as a percentage of the initial wounded area.

### Tube formation assay

Tube formation was investigated as reported previously with some modifications [23]. 96 well plates were coated with Matrigel™ (Corning, USA) and overlaid with 2 x 10^4^ serum-starved HUVECs per well. Each triplicate wells were incubated with the same treatment of non-DM and DM-MSC CCM, complete M199 and SFM at 37°C with 5% CO_2_ for 12 hours. Tube formation was evaluated using an inverted microscope and images were analyzed by Wimasis software.

### Statistical analysis

For each treatment, 3 biological samples were performed and each sample was analyzed in triplicate. The data are presented as the means ± SD, and analysis of variance (ANOVA) was applied for the statistical analysis. The statistical analyses were performed using the program Graph Pad Prism 5.0. Differences were considered statistically significant at *P-value* < 0.05.

## Results

### Comparison of non-DM and DM-MSCs

#### Morphology

The adherent MSCs of non-DM and DM subjects grew in a spindle shape which is a typical fibroblast-like cell morphology (Figure 1). However, at early passages DM-MSCs were longer and flatter (Figure 1, A) compared to non-DM MSCs (Figure 1, B). At later passages the DM-MSCs had enlarged individual cell area, flat shape with short blunt edge projections (Figure 1, C) compared to non-DM MSCs (Figure 1, D).

**Figure 1:**
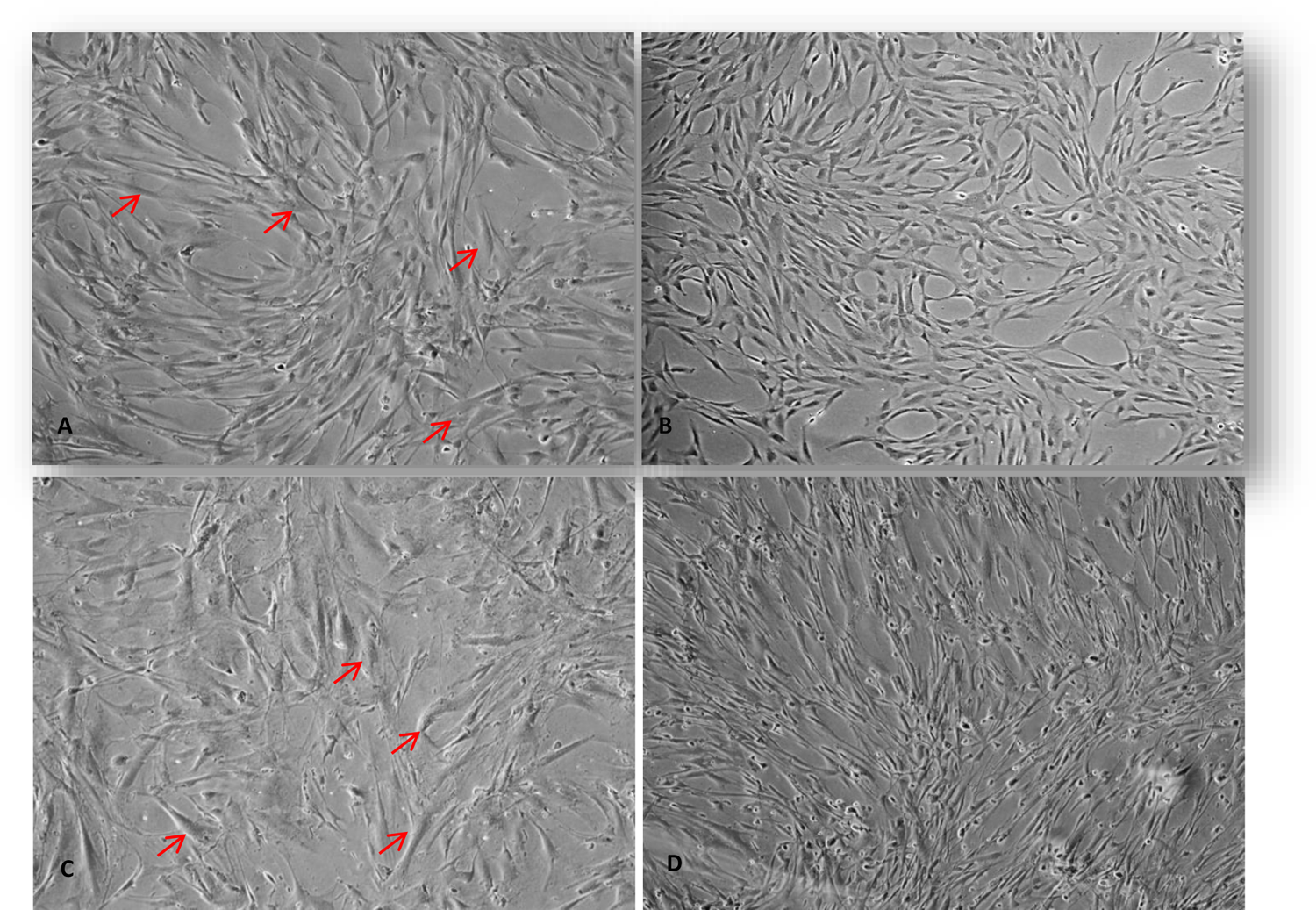
Morphology of MSCs at early and late passages. A: DM MSCs at passage 4. B: non-DM MSCs at passage 4. C: DM MSCs at passage 13. D: non-DM MSCs at passage 13. Arrows point to the flat and enlarged morphology of DM-MSCs appeared in early and late passages (Magnification 10X).

#### Growth kinetic assessment

Population doubling time (PDT) was measured from passage 4-12 for the non-DM and DM MSCs. Results in Figure 2 show no significant difference in the proliferation potential of non-DM MSCs from passage 4 to passage 10 compared to DM-MSCs, nonetheless the proliferative potential of non-DM MSCs was significantly higher for P10-P11 and P11-P12 compared to DM MSCs (20 ± 1.6; 23±0.8 and 38 ± 0; 32±1.7 hours) respectively (*p-value* < 0.05).

**Figure 2:**
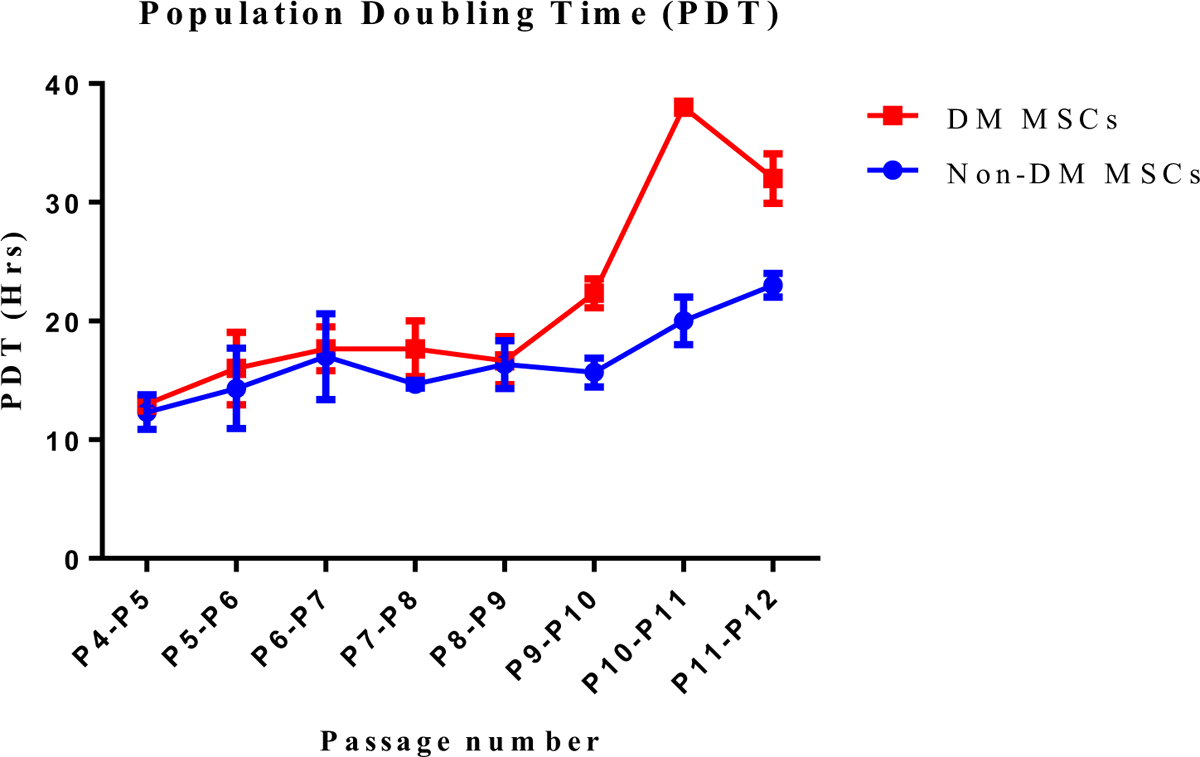
Population doubling time (PDT) for non-DM (circles) and DM-MSCs (squares) for passages 4-12. Data expressed as mean ± SEM (n=3). *(*p-value* < 0.05).

#### Immunophenotypic characterization

Purified BM-MSCs could be easily characterized by cell markers expressed on their surface. The expression of MSC surface markers by flow cytometer showed similar expression pattern in non-DM and DM-MSCs (Figure 3).

**Figure 3:**
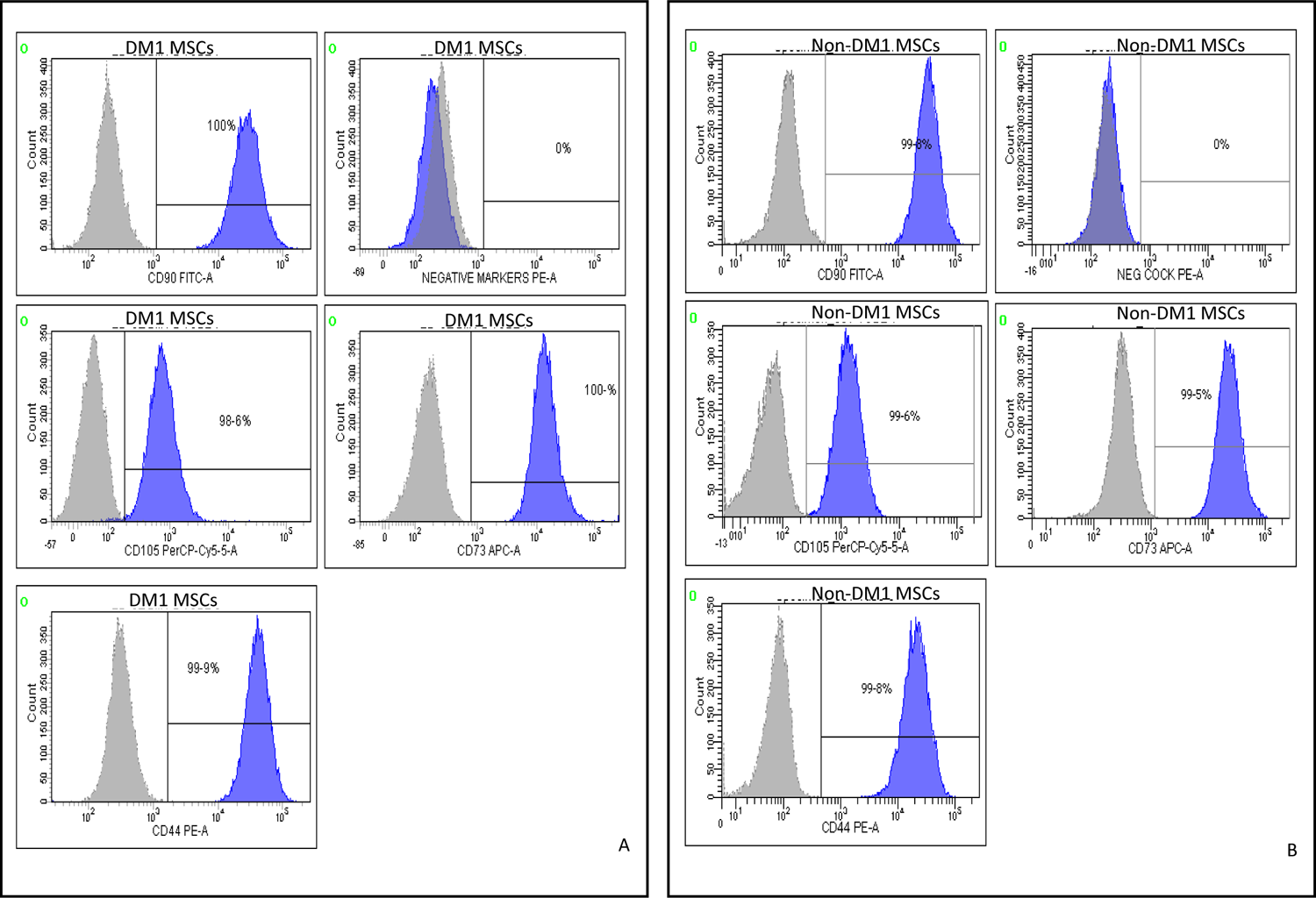
Flow cytometry analysis of non-DM and DM MSC surface marker expression. The data shown are representative cell phenotype analyzed at passage 4. Gray peaks correspond to the isotype control and the blue peaks to the antibody of interest. (A) DM-MSCs (B) non-DM MSCs.

#### Multilineage differentiation potential

For both differentiation protocols, the non-DM and DM MSCs treated as controls were cultivated with expansion medium during the entire protocol, as expected, showed no formation of lipid vacuoles or calcium deposits (Figure 4, A and B and Figure 5, A and B). Adipogenic differentiation was confirmed by the accumulation of lipid vacuoles and oil red O staining. Higher lipid accumulation and more differentiated MSCs were observed at day 21 of non-DM MSCs differentiation (Figure 4. C&D). Osteogenic differentiation was demonstrated by matrix calcification and mineral deposition detected through Alizarin red S staining. After 21 days of induction, the osteogenic differentiation capacity detected in DM-MSCs was greater compared to non-DM MSCs. (Figure 5 C and D).

**Figure 4:**
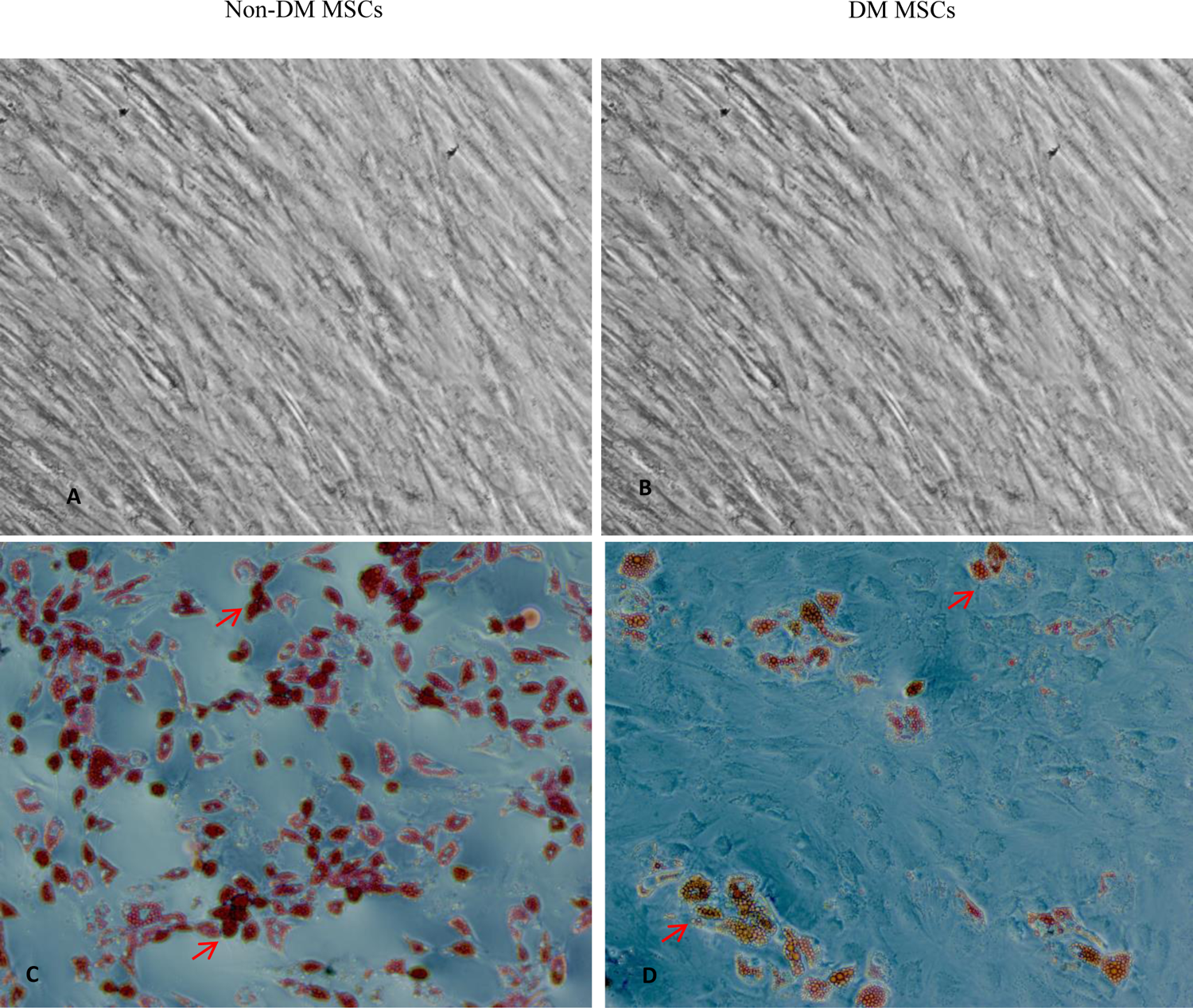
Representative sample of BM-MSCs adipogenic differentiation. After 21 days of induction, lipid droplets were stained with oil red O. A&B undifferentiated cells. C&D stained differentiated cells (Magnification 10X).

**Figure 5:**
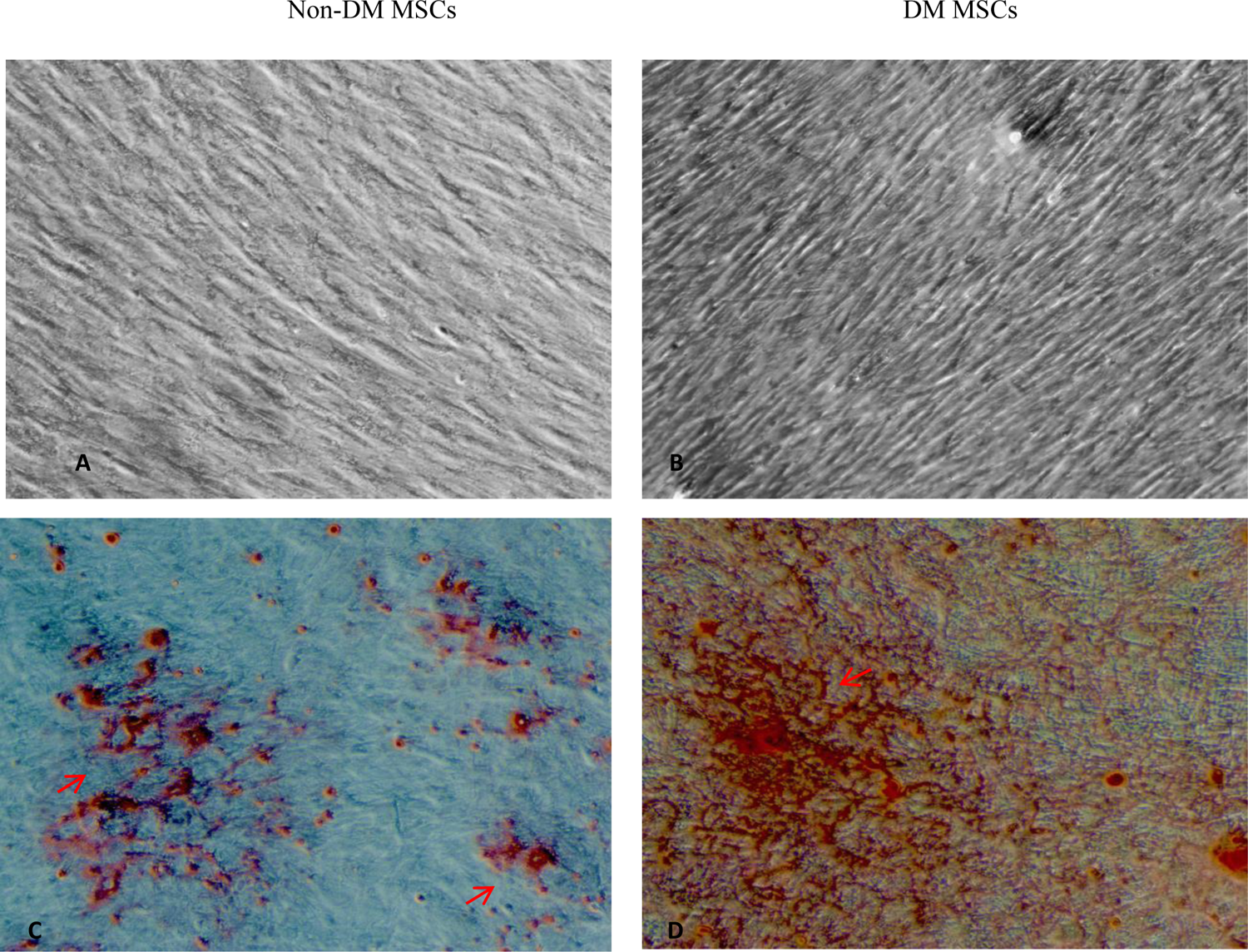
Representative sample of BM-MSCs osteogenic differentiation. After 21 days of induction calcium mineralization was visualized by Alizarin red stain; A and B are undifferentiated cells. C and D are differentiated cells stained with Alizarin red S staining. (Magnification 10X).

#### Angiogenesis proteins

A semi-quantitative test was performed to evaluate the secreted angiogenesis-related proteins in the conditioned medium. In order to exclude any analyte that was already in the medium needed for the maintenance of HUVECs, SFM was used as a control. Results show that SFM medium consisted of four angiogenesis-related proteins; serpin-E1, serpin-F1, thrombospondin-1, and TIMP-1 (Figure 6, A). Nonetheless, when these four proteins were excluded, 9 and 10 proteins were found in non-DM and DM-MSC CCM, respectively (Figure 6, B and C). The protein expression levels were measured by the Image Studio Lite; 5.2 software and presented as the mean pixel density normalized by the positive spot reference (Figure 7). Activin A was unique in Non-DM MSC CCM whereas angiopoeitin-1 and IGFBP-2 were only found in DM MSC CCM. When the relative concentration of TIMP-1, thrombospondin-1, serpin-F1, and serpin-E1 proteins of SFM were compared to CCM, only thrombospondin-1 was found to be over-expressed in the CCM secretome compared to SFM. Whereas serpin-E1, serpin-F1, TIMP-1 proteins were down-regulated in CCM of non-DM and DM MSC (2.6-, 3.7- and 2.0-fold change; 2.2-, 6.2-, and 14.2-fold change), respectively (Supplemetary Table S1). In addition, we found that the relative protein expression levels of uPA, IGFBP-3, and angiogenin were upregulated in DM-MSC (1.1-, 1.3-, and 1.5-fold change, respectively). On the contrary, five angiogenesis proteins were clearly upregulated in non-DM MSC secretome: VEGF, PTX3, PF4, MCP-1 and FGF-7 (1.7-, 4.2-,1.5-, 1.0-, 2.4-fold change), respectively (Supplementary Table S2). Nonetheless, no significant difference was found between the common analytes in the conditioned medium.

**Figure 6:**
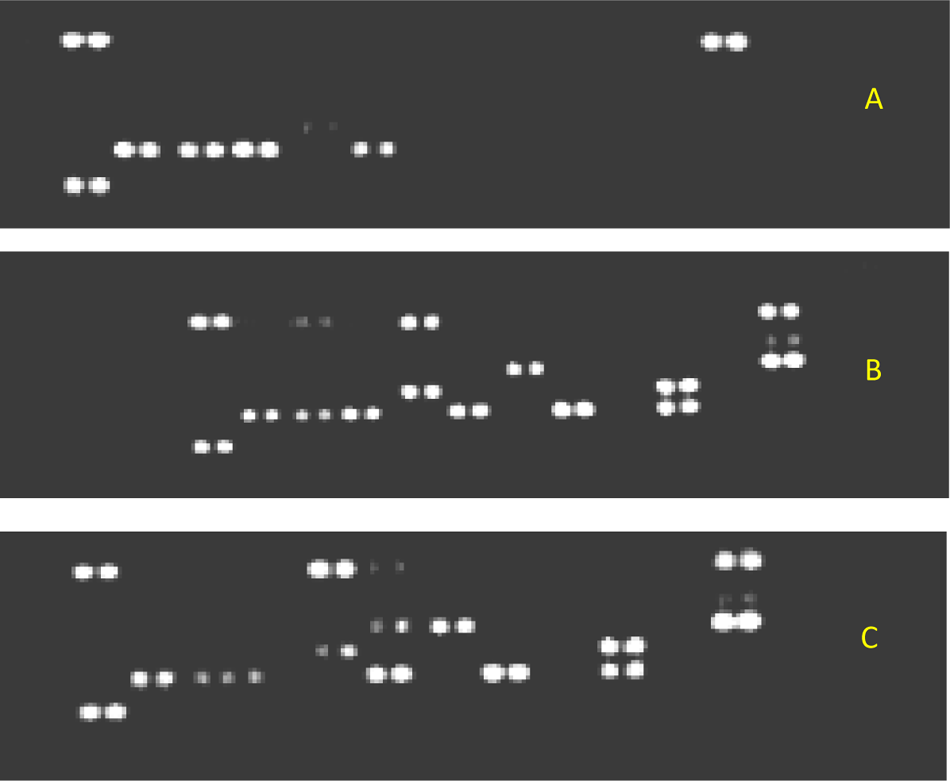
Membrane image after exposure to X-ray film. Human angiogenesis array membranes were used to analyze secretion of angiogenic growth factors of (A) SFM (B) non-DM MSC CCM (C) DM MSC CCM.

**Figure 7:**
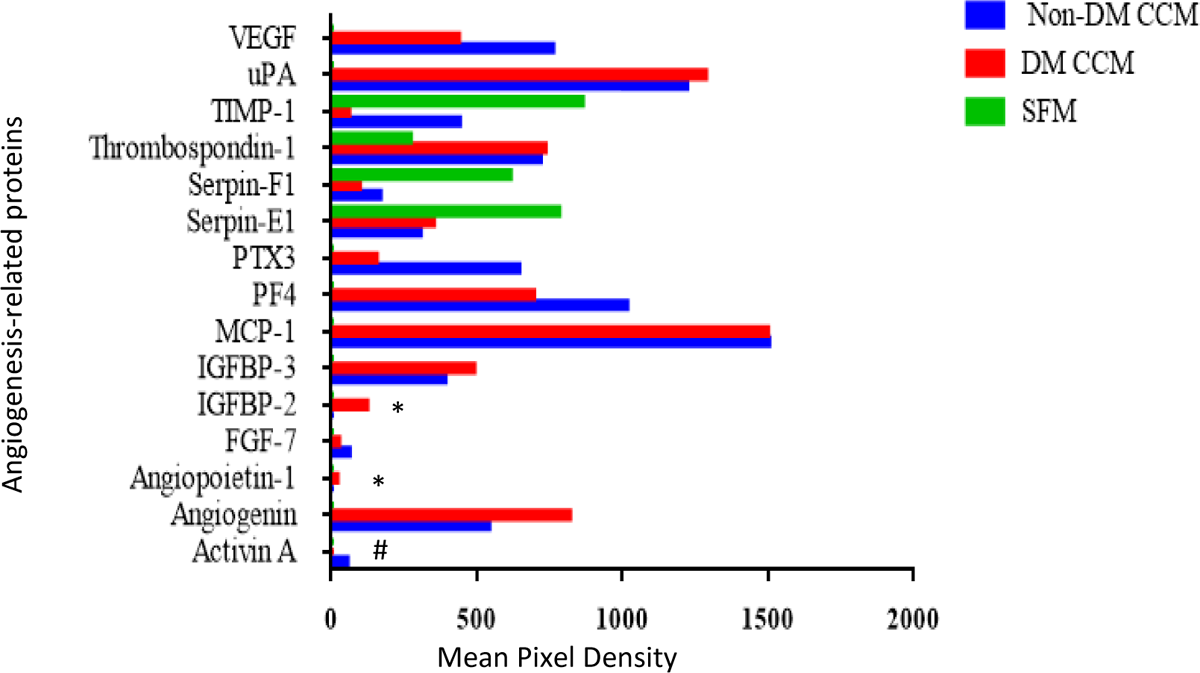
Mean pixel density of angiogenesis-related proteins detected in conditioned medium. The relative protein expression levels of levels of 15 angiogenesis-related cytokines detected in the three test groups were measured by the Image studio software and presented as the mean pixel density normalized by the positive control. * detected only in DM MSC CCM, # detected only in non-DM MSC CCM.

### Isolation and characterization of HUVEC

#### Characterization of HUVEC in vitro

Primary HUVECs became attached to the gelatinized surface after 24 hours and became confluent and ready for subculture after 5 days. Cells grew in the form of island-like colonies of homogeneous polygonal cells with distinct cell boundaries (Figure 8, A). After P_0_, confluent cells had polygonal cobblestone appearance (Figure 8, B). Isolated HUVECs were analyzed for cell surface expression for CD45, CD34, CD144 and CD31. HUVECs were found to express mature endothelial cell markers CD144 and CD31 and negative for CD34 (Supplementary Figure S3).

**Figure 8:**
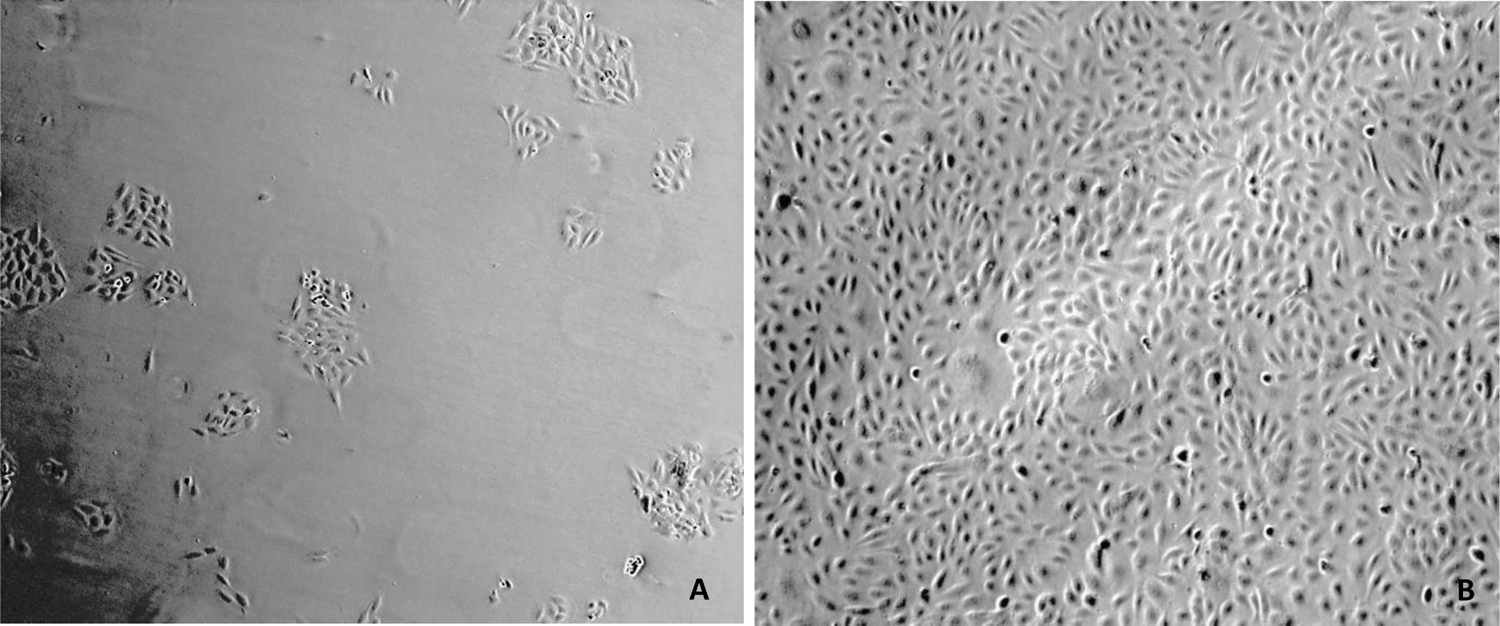
Morphology of HUVECs at P_0_ after 24 hours of isolation (A), confluent HUVECs at P_1_ (B). (Magnification 10 X)

#### The effect of MSC-CCM on cell viability

The viability of cells grown in non-DM and DM-MSC CCM was found to be significantly increased as compared to both controls (*p* < 0.05), however there was no significant difference between cells grown in non-DM and DM-MSC CCM (Figure 9).

**Figure 9:**
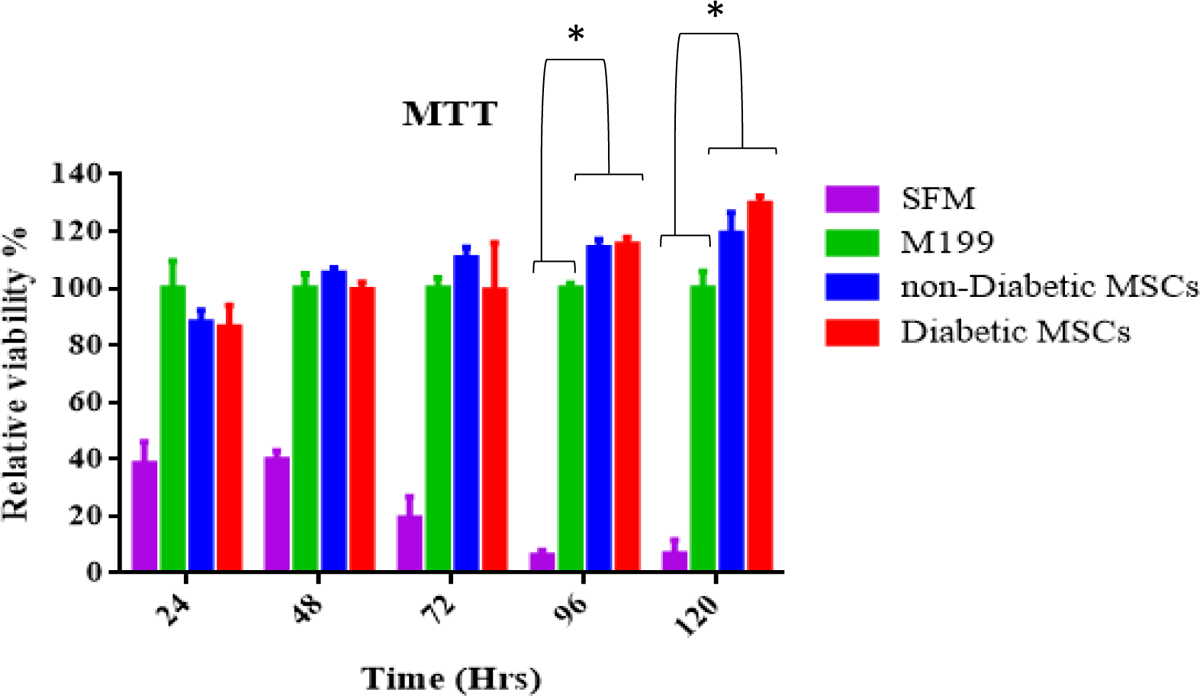
MTT assay performed at different time points. At 96 and 120 hrs, CCM from non-DM and DM-MSC significantly enhanced the viability of HUVECs as compared to M199 and SFM (*p value* < 0.05). Results are represented as mean ± SD of three independent experiments.

#### Assessment of cell migration of HUVECs using transwell membrane

The number of migratory cells was quantified by counting 10 random fields of each replica (Figure 10). There was a significant increase in the number of ECs migrating toward the CCM of both non-DM and DM groups in comparison to the SFM (*p* < 0.05). However, number of migratory HUVECs toward M199 (124.4 ± 13.3) was significantly greater than cells migrating toward CCM from non-DM and DM MSCs (72.8 ± 1.7, 59.0 ± 4.0) respectively (*p* < 0.05). Nonetheless, there was no significant difference in migration of HUVECs toward the non-DM-CCM compared to DM-CCM (*p* > 0.05).

**Figure 10:**
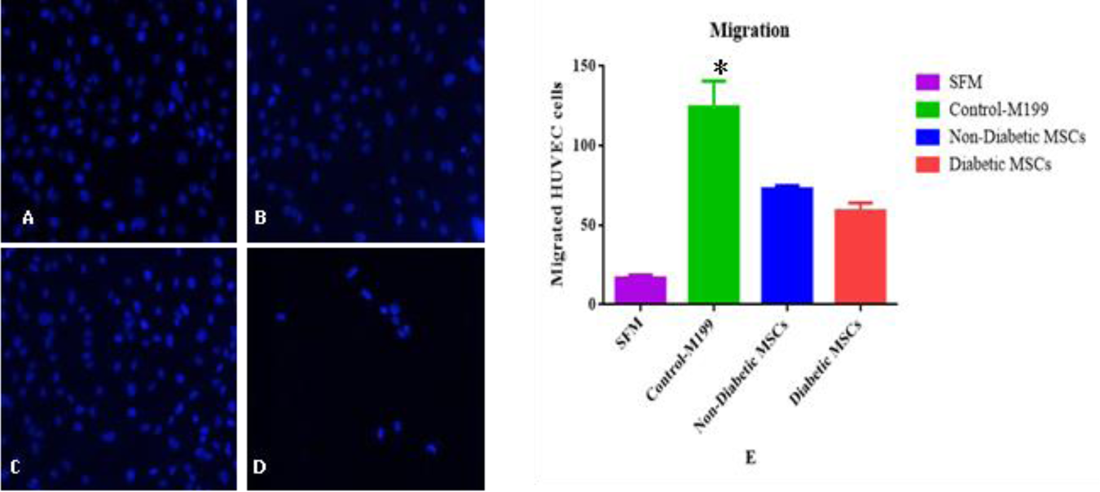
Representative fields of transwell migration of HUVECs among different treatments A: non-DM-, B: DM-MSC CCM, C: M199, D: SFM. E: Quantitative analyses of transwell migration of HUVECs among different treatments. Migration enhancing effect is maximal with M199. Data presented as Mean ± SD for each condition in triplicate wells counted in 10 random fields.

#### Wound Healing

The extent of wound healing was observed with regard to cell conditioned media added and images were taken at the indicated periods (Figure 11, A). The percentage of the remaining open area compared to the initial wound area was quantified (Figure 11, B). The scratch wound healing revealed that compared with non-DM CCM treated HUVECs, wound healing capacity of DM CCM treated HUVECs was attenuated, (25.3% ± 6.3, 49.3% ± 10.1 respectively) but it was statistically insignificant (*p-value* >0.05) (Figure 11).

**Figure 11:**
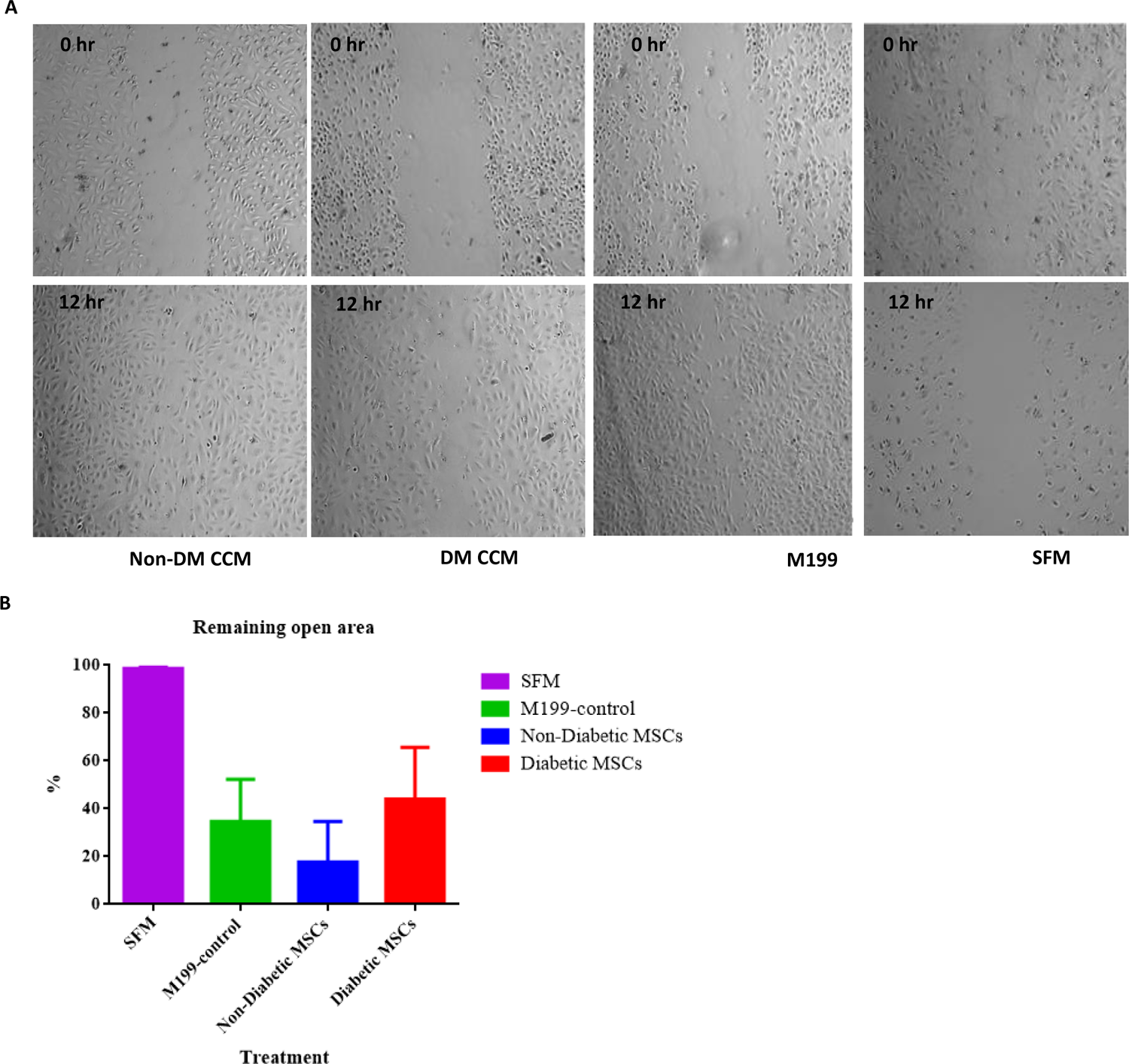
(A) Closure of an artificial wound in HUVEC monolayer at 0 and 12 hours incubated with CCM, M199 and SFM (magnification 4 X). (B) The percentage of remaining open area after 12 hours. Data presented here as mean ± SD for each condition in triplicate wells. *p-value* >0.05

#### Tube formation assay

Images were documented through inverted phase contrast microscopy for 12 hours to test the *in vitro* differentiation of HUVECs (Figure 12). Several characteristics were used to quantify tube network formation. Number of nodes and mean tube length were significantly higher in non-DM CCM group compared to controls (*p* < 0.05) but no significant differences were measured between non-DM and DM MSC CCM groups (*p* > 0.05) (Figure 13). Cell-conditioned medium derived from both non-DM and DM MSCs enhanced the formation of HUVEC tubular networks as early as 4 hrs following seeding onto the matrigel matrix (Figure 12). No significant change was found in total number of tubes formed by HUVEC cultured in DM-MSC CCM (31.7± 2.5) compared to cells cultured in non-DM CCM (44 ± 3.7) per field (Figure 13, A) (*p* > 0.05). Non-DM MSC CCM was efficient at stimulating the growth of HUVEC tubular structures, this was demonstrated through quantification of mean tube length (Figure 13, B) and total number of nodes (*p* < 0.05) (Figure 13, C).

**Figure 12:**
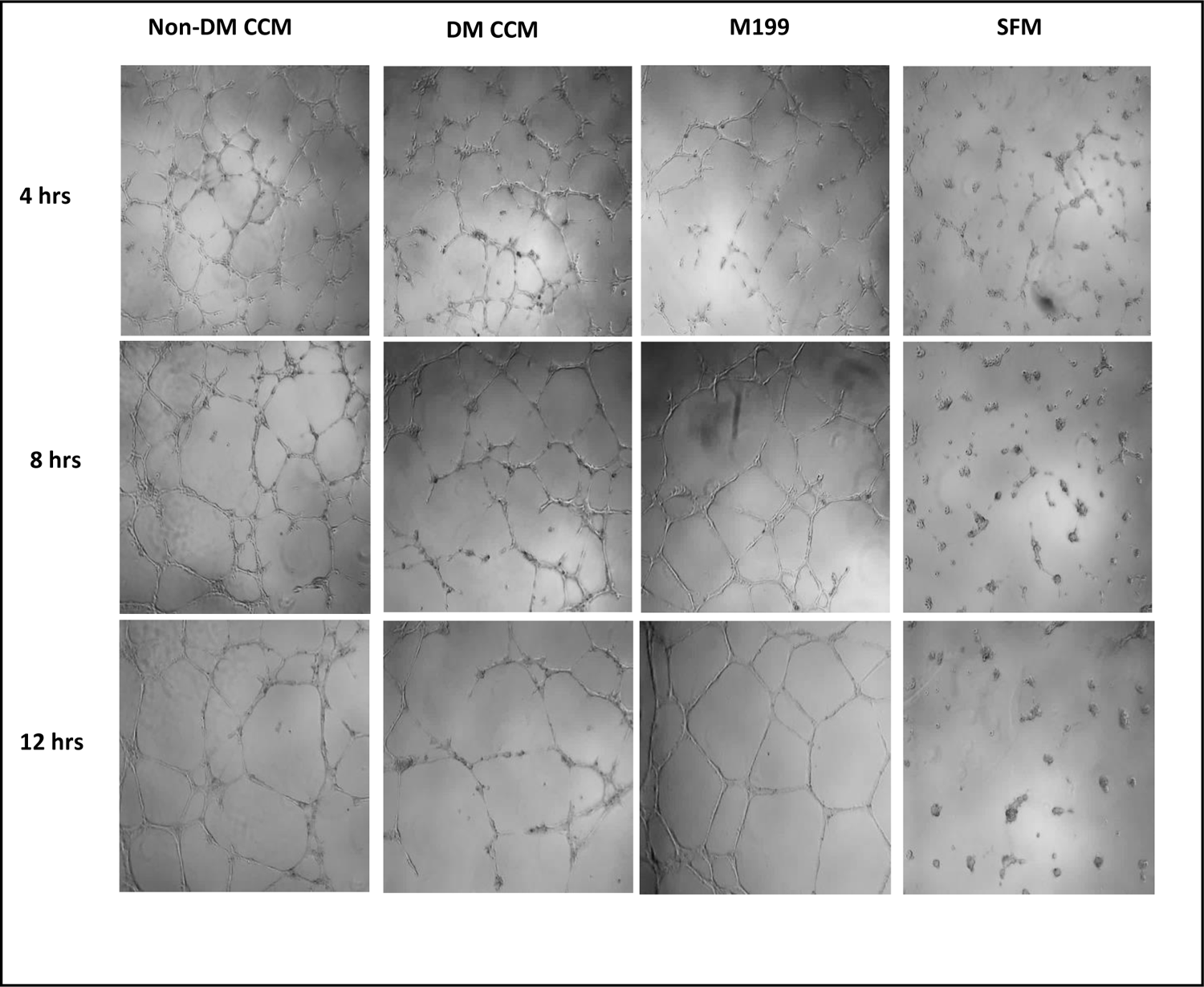
Representative images of capillary network formation captured at 4, 8 and 12 hrs. HUVECs cultured on Matrigel coated plates treated with non-DM MSC CCM, DM-MSC CCM and M199 assemble into tubular structures. (Magnification 10X)

**Figure 13:**
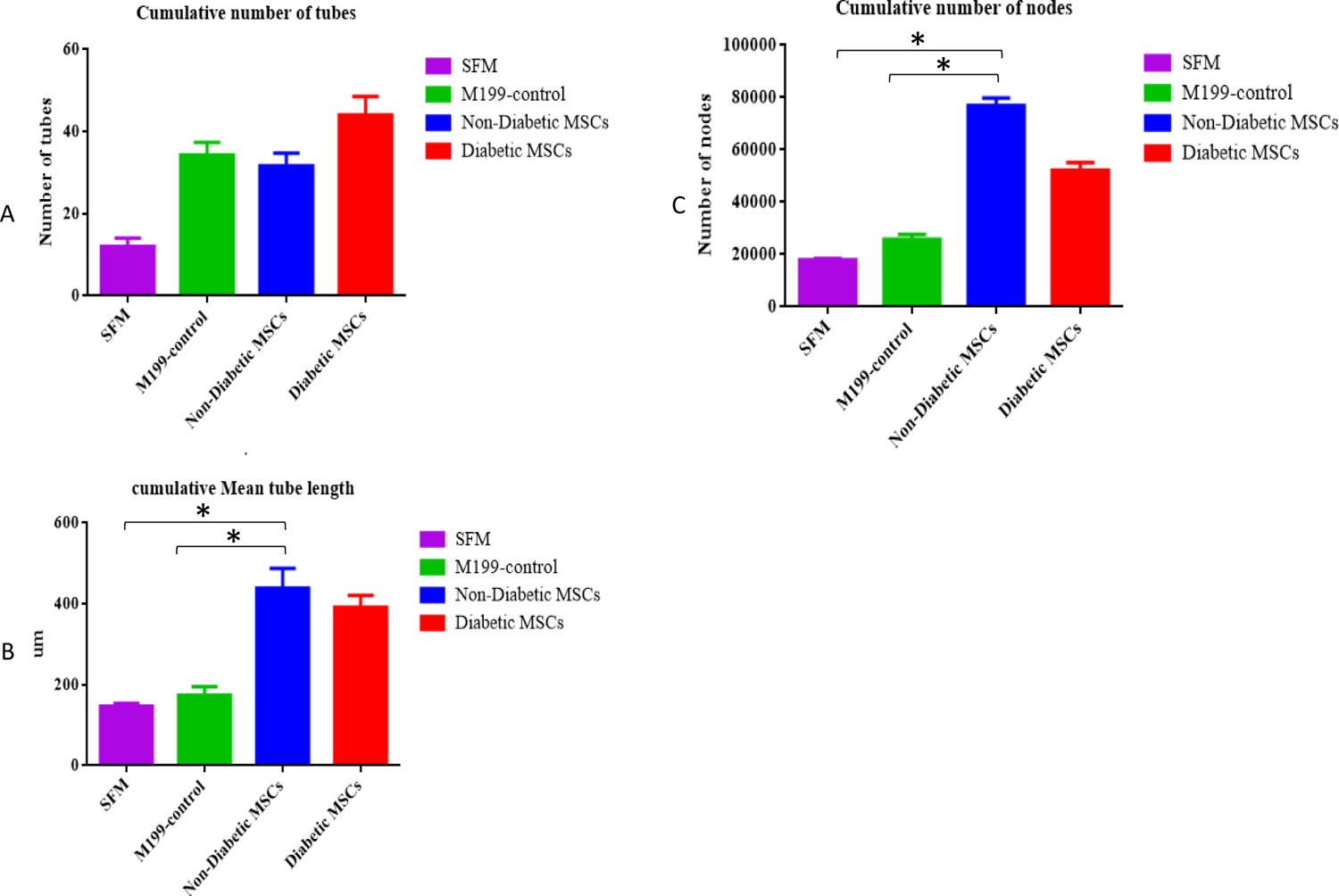
Effect of CCM derived from non-DM and DM-MSCs along with M199 (positive control) and SFM (negative control). Analysis of cumulative (A) number of tubes (B) mean tube length and (C) number of nodes during the 12-hour incubation. Data presented here as mean ± SD for each condition in triplicate wells/field.f

## Discussion

MSCs and their potential clinical applications are attracting major attention. Therefore, it is essential to maximize their therapeutic benefits by optimizing the function and the proliferative capacity of MSCs. Despite their beneficial effects, stem cells have provided short-lived benefit in many human clinical trials [24]. It is suggested that the underlying cause of this disappointing point is the fact that significant number of transplanted cells fail to survive and proliferate enough to exert profitable effects [25]. The cause of the poor stem cell survival after transplantation is not fully understood yet. Therefore, stem cell research needs to be more oriented toward a better understanding of the best conditions that will enhance their survival and expansion in order to boost their therapeutic capacity following transplantation. Hyperglycemia that is encountered in many pathological diseases such as DM, was reported to cause dysfunction in bone marrow hematopoietic niche [10]. DM also negatively affects the mobilization and functions of adult stem cells [26]. Furthermore, high glucose induced morphological changes and premature BM-MSC senescence in culture medium [11,27]. In order to optimize the therapeutic potential of autologous MSC in diabetes vascular complications. Consistent with our results, a study conducted on adipose derived MSCs (AD-MSCs) by a research group demonstrated that the majority of AD-MSCs isolated from diabetic patients exhibited a cocktail of morphologies including flat spread-out cells of irregular shape [28]. This finding was also supported by Stolzing *et al.* (2010) who observed that BM-MSCs become less confluent and that the accumulation of enlarged cells was elevated in a time-dependent manner when exposed to diabetic environment [29]. Growth kinetics of our MSCs represented by population doubling time showed higher proliferation potential of non-DM MSCs at higher passages. It has been also elucidated that the proliferation of the diabetic AD-MSCs was slower than that of lean AD-MSCs, which was strongly reduced by the end of 120 hours in culture [28]. Interestingly, Coutu *et al.* (2011) depicted that morphological alterations and reduced proliferative activity of diabetic AD-MSCs were recovered after the exposition of these cells with β-FGF [30]. β-FGF promotes MSC proliferation by arresting cellular senescence through Akt signalling pathway; thereby, it can be a key component in the maintenance of MSCs’ self-renewal and stemness. Therefore, supplementation of the culture medium of DM-MSCs with β-FGF could be considered as a conditioning strategy for DM-MSCs to rescue cells from reduced proliferative capacity and senescence. Our results indicated that non-DM MSCs possess greater adipogenic differentiation potential compared to DM MSCs. This was evident from the earlier adipocyte differentiation and higher number of differentiated adipocytes of the non-DM MSCs, while a greater osteogenic differentiation potential was detected in DM-MSCs. Ezquer *et al.*, (2011) reported that viable MSCs obtained from diabetic mice have the same adipogenic potential than that of normal mice [31]. They also reported that osteogenic potential of diabetic MSCs differs from healthy ones. Correspondingly, it has been shown that the osteogenic differentiation potential of diabetic MSCs was significantly reduced compared with that obtained from age matched healthy MSCs [29]. Opposed to this data, it was observed a variability in the adipogenic differentiation potential of BM and gestational MSCs, which was greater in MSCs cultured in high glucose [32].

Nonetheless, the latter study revealed similarity in the osteogenic differentiation potential of healthy and diabetic MSCs. Such contradicting results might be due to differences in the isolation, culturing, and differentiation protocols. In addition, longer differentiation period might be required by one cell type to achieve a similar level of differentiation [33]. The focus of our study was also to further investigate the impact of the diabetic milieu on the composition of BM-MSC secretome, and hence, its angiogenic potential on ECs. The aim of targeting this point was to illustrate the extent to which MSCs can have beneficial contribution in the treatment of diabetic vascular complications. Therefore, developing relevant aspects for future BM-MSC based therapies to enhance diabetic tissue repair. The relative concentration of the secretome components was assessed; we demonstrated that Activin A was exclusively secreted by non-DM MSC. Activin A which is a transforming growth factor (TGF)-β superfamily member, was found to be secreted from mesenchymal stem cells isolated from bone marrow, tonsils, muscles, and dental pulps [34]. It is a pivotal regulator in the maintenance of pluripotency of stem cells by upregulating the expression of the key embryonic transcription factors Oct4, Nanog, and Sox2 [35]. In 2014, Chatterjee *et al*. reported the role of Activin A produced by MSCs in modulating NK cells function by suppressing their IFN-γ production. However, the role of Activin A in angiogenesis is controversial [36]. While it was demonstrated to inhibit the cellular growth of vascular endothelial cells [37–39], contrasting findings showed that Activin A induced vascular endothelial cells proliferation [40]. Supporting this observation, Activin A was also found to induce VEGF expression leading to recruitment of blood vessels *in vitro* [41].This discrepancy of the pro- and anti-angiogenic role of activin A can be explained by the dual role of TGF-β [39]which share the Smad intracellular signalling proteins with TGF-β [42]. It has been postulated that the local concentration of activin A may dictate whether it has a pro- or anti-angiogenic effect [43]. Vascular endothelial growth factor (VEGF) and angiopoietin-1 are key regulators of angiogenesis and vascular maturation [44]. VEGF is a one of the most essential proangiogenic growth factor, which directs vascular outgrowth of tip cells by sensing VEGF gradient [45]. It is also evident that VEGF is active during wound healing; its levels affect the speed and quality of healing. Insufficient levels of VEGF impair the repair process and develop chronic and nonhealing wounds [46]. Supernatant derived from non DM-MSCs showed an elevated concentration of VEGF, nonetheless, it was also found in DM-MSC derived supernatant. This reflected on the enhanced sprouting of our ECs when cultured in both non-DM and DM MSC CCM. Same observation was described by Beckermann *et al.* in which the secretion of VEGF by MSCs under basal conditions increased sprouting, supporting angiogenic potential of MSCs [47]. Monocyte chemotactic protein-1 (MCP-1) is recognized as a proangiogenic chemokine, it is reported to induce angiogenesis through the up-regulation of HIF-1α and subsequent induction of VEGF [48]. MCP-1 was found to be elevated in our CCM of non-DM as well as DM MSCs. In agreement with Ribot *et al.*, IGF-1 was reported to be up-regulated in DM-MSCs [49]. IGF-1 has been shown to be modulated by IGFBP-2 and it is a critical regulator of angiogenesis [50]. However, IGFBP-2 and Angiopoietin-1 were absent in our non-DM MSC CCM. This could be as a result of variations in the degree of confluence at the time of CCM preparation, giving rise to a varying number of MSC in culture hence different concentration of our protein components. It is also noteworthy to mention that paracrine effects of MSCs might not support the initiation of the angiogenic process, but rather contribute to guiding the sprouting blood vessels [51]. Whereas others reported the enhanced proliferation of EC by BM-MSC [52]. Albeit the variability of the MSC supernatant composition, several pivotal angiogenic factors were found in both non-DM and DM MSCs and their relative concentrations were somehow close. However, to get a deep perception of diabetes impact on MSC secretome, a quantitative proteome analysis along with molecular testing should be performed. Despite the contribution of BM-MSC paracrine factors in neovascularization and tissue repair, the effect of DM on the BM-MSC secretome and its impact on angiogenesis are still vague [49,52,53]. There are debatable results on the angiogenic potential of DM-MSC. A group of researchers demonstrated that the neovascularization and *in vivo* therapeutic effects for hindlimb ischemia recovery of MSCs derived from diabetic subjects were defected [12]. In line with these results, bone marrow mononuclear cells from diabetic mice exhibited a reduced ischemia-improving potential [54]. While another study addressed the multipotency defect and impairment of angiogenic differentiation potential in DM-MSCs, resulting in reduced blood flow recovery after limb ischemia [55]. A similar observation was found by Dzhoyashvili *et al.* where they reported that secretome derived from AD-MSC exhibited a declined angiogenic potential [56]. Contrasting findings in one research provided evidence that the secretome composition of DM-MSCs differs from non-DM MSCs and enhances angiogenic capabilities. The results of this study showed that several pro-angiogenic genes were overexpressed whereas anti-angiogenic genes were downregulated. Another major finding of this study is that the DM-MSC CCM promoted *in vitro* endothelial cell migration and tubular structure formation, along with *in vivo* vascular formation [49]. A correlative response was observed for BM-MSCs cultured under *in vitro* hypoxic and hyperglycemic conditions to simulate *in vivo* diabetic environment [57,58]. This suggests that MSCs experiencing simulated major insults of DM can survive and remain functional [57]. Concordant to these observations, one study showed that DM-MSCs were also capable of enhancing the metabolic profile of diabetic animal model, highlighting the therapeutic potential of DM-MSCs [59]. Furthermore, a previous study from the same group investigated the DM-MSCs potential in rescuing cardiac electric changes caused by diabetes [60]. Our study showed that BM-MSCs isolated from both diabetic and non-diabetic subjects enhanced the viability of endothelial cells as seen in MTT assay. Furthermore, both BM-MSC groups boosted the motility, migration and tubular structural formation of HUVECs. However, the difference in the effect of both BM-MSC groups on the angiogenic properties of ECs were not statistically significant. Consequently, disturbed balance between pro- and anti-angiogenic factors may give rise to perturbations in angiogenesis, in which, excessive angiogenesis occurs in diabetic retinopathy and nephropathy, whereas reduced angiogenesis participates to diabetic impaired wound healing [61]. This emphasizes the paracrine effect of an altered BM-MSCs CCM on endothelial cells. In addition, it highlights the dualistic contribution of angiogenesis to the pathogenesis of diabetic vascular complications.

Our study showed that BM-MSCs isolated from both diabetic and non-diabetic subjects enhanced the viability of endothelial cells as seen in MTT assay. Furthermore, both BM-MSC groups boosted the motility, migration and tubular structural formation of HUVECs. However, the difference in the effect of both BM-MSC groups on the angiogenic properties of ECs were not statistically significant.

Despite the multiple *in vitro* divisions of DM-MSCs in a normoglycemic conditions, the impact of the diabetic milieu on the MSC behavior is still evident as seen in the aforementioned studies [10,26,28]. This can be corroborated by the “metabolic memory” phenomenon. This phenomenon proposes that the diabetic environment can lead to dysregulation in epigenetic mechanisms altering the expression of pathological genes in target cells without changes in the consecutive DNA sequence [62].

Last, but not least, this argumentative data cannot change the fact that the impact of MSCs on patients with peripheral vascular diseases hold great promise [63]. Yet, it is crucial to emphasize another fact that diabetic populations are on the rise and these patients are at high risk of cardiovascular diseases [64]. As these individuals are most likely targeted for cell therapy, it is essential to test MSCs isolated from such subjects and define the limitations in their therapeutic potential particularly vessel formation capabilities. Still different approaches and mechanisms should be proposed in this field to restore the defective characteristics of MSCs derived from diabetic subjects, with differences in isolation methods, species and endothelial cell sources should be taken into account [65].

## Conclusion

In general, we can conclude that despite the negative impact of the hyperglycemic milieu on the proliferation potential of the MSCs, the angiogenic-related functional properties of MSCs remain intact and they are able to survive the harsh diabetic conditions. This is essential for clinical applications since the number of diabetic patients is increasing and they are the most targeted population in MSC-based clinical approaches.

